# Internalizing cooperative norms in group-structured populations

**DOI:** 10.1101/722439

**Authors:** Erol Akçay, Jeremy Van Cleve

## Abstract

Social norms play a crucial role in human behavior, especially in maintaining cooperation within human social groups. Social norms might be self-enforcing or be enforced by the threat of punishment. In many cases, however, social norms are internalized and individuals have intrinsic motivations to observe norms. Here, we present a model for how intrinsic preferences to adhere to cooperative norms can evolve with and without external enforcement of compliance. Using the methodology of preference evolution, we model how cooperative norms coevolve with the intrinsic motivations to follow them. We model intrinsic motivations as being provided by guilt, a kind of internal “punishment” that individuals feel for falling short of cooperative norms, and show that the shape of this internal punishment function plays a crucial role in determining whether and how much internalization can evolve. We find that internal punishment functions that eventually level off with the deviation from the norm can support internalization without external punishment. In contrast, internal punishment functions that keep escalating with the deviation from the norm require external punishment, but yield stronger norms and more cooperation when external punishment is present. By showing how different preference mechanisms can enhance or limit norms that stabilize cooperation, these results provide insights into how our species might have evolved the norm psychology that helps us maintain such complex social and cultural institutions.

## Introduction

The success of humans in spreading through all of Earth’s ecosystems and transforming them at planetary scale is directly dependent on our capacity to cooperate in large groups and self-organize in complex social structures that sustain such cooperation. One of the main components of such large-scale cooperation is the human capacity and propensity for inventing and following social norms (Ostrom 2000; Fehr and Schurtenberger 2018). Social norms influence almost all aspects of human behavior, providing a “grammar of society” (Bicchieri 2005, 2010) that constrains and enables different kinds of individual behaviors, coordinates collective behavior, and can sustain cooperation in the face of conflicts of interests.

Although commonly discussed as a single phenomenon, social norms are best seen as a diverse set of emergent phenomena that result from the interaction of mechanisms at multiple scales, from individual level cognition to population level gene-culture coevolution (Gintis 2003). Some social norms turn social dilemmas into coordination games (Bicchieri 2005, 2010) by prescribing particular behaviors and inducing individuals to expect others to behave the same way while other norms are signals that coordinate individual behavior to implement outcomes (i.e., correlated equilibria) that improve on the outcomes possible without such signals (i.e., Nash equilibria) (Gintis 2010; Morsky and Akçay 2019).

Prescriptive norms will have little impact on actual behavior unless there are some mechanisms that enforce them. Some norms are self-enforcing in the sense that once they are established, it is in the self-interest of agents to follow them (Binmore 1998). Other social norms, however, may require the threat of institutional or peer punishment to make individuals adhere to them (Gintis 2003). It is well-established that by sufficiently punishing individuals who deviate from the norm one can enforce a wide range of outcomes in social dilemmas (Ostrom, Walker, and Gardner 1992; Boyd and Richerson 1992; Fehr and Gachter 2000; Gintis 2000; Henrich et al. 2010). However, since costly punishment is itself a public good whose provision requires overcoming another social dilemma (the “second-order free-rider problem” Heckathorn 1989), explaining the evolution of social norms also requires explaining the evolution of the mechanisms that sustain them.

Finally, many social norms are followed by individuals because they are internalized; in other words, individuals have acquired intrinsic preferences to comply with norms even if such compliance is costly for their material interests. Internalization of norms is a long- and widely-recognized fact of human social life (Chudek and Henrich 2011). Intuitively, we are all familiar with the concept: we follow countless norms daily, at varying inconvenience to ourselves, even when we run little risk of detection or punishment for not complying. Experimental evidence suggests that people’s behavior in laboratory games are affected by their beliefs about other’s expectations (Dufwenberg, Gächter, and Hennig-Schmidt 2011) and by variation in their sensitivity to norms (Kimbrough and Vostroknutov 2016). Such intrinsic preferences for norm compliance may be modulated by particular neural circuitry in the brain (Spitzer et al. 2007; Ty, Mitchell, and Finger 2017). Theoretical accounts of intrinsically motivated compliance with social norms have modeled how parents might invest into socializing their offspring to internalize different preferences (Bisin and Verdier 2001), how guilt from failing to live up to others’ expectations can drive individual behavior (Battigalli and Dufwenberg 2007), and how natural or cultural selection might favor intrinsic preferences for norm compliance in social interactions (Gintis 2003; Gavrilets and Richerson 2017). Our chapter contributes to this literature by modeling the coevolution of social norms and their internalization.

Social decision making involves cognitive mechanisms that evaluate the direct benefits and costs of potential social behaviors. In the context of norms, these benefits and costs will include how different behaviors compare with the norm. A simple way to summarize how these cognitive mechanisms might work is to assume that social behaviors generate internal (i.e., neurophysiological) signals of reward or punishment, and that individuals behave in such a way as to increase their internal reward signals and decrease internal punishment signals. For example, in the presence of a contribution norm to a public good, an internalized norm (or beliefs of others’ expectations) can induce a subjective reward for complying with the norm or subjective displeasure for falling short (Axelrod 1986; Bicchieri 2005; Dufwenberg, Gächter, and Hennig-Schmidt 2011; Kimbrough and Vostroknutov 2016). These internal rewards or punishments (e.g., feelings of guilt) can create instrinsic motivations to follow norms and reduce or obviate the need for external punishment or reward.

The internal signals driving decision-making are sometimes called “preferences” and their evolution can be studied using mathematical models (Güth 1995; Dekel, Ely, and Yilankaya 2007; Akçay et al. 2009; Alger and Weibull 2013). In these models, individuals have genetically or culturally transmitted traits that determine their preferences, represented by a utility or objective function. This function typically depends on individuals’ material payoffs but need not be identical to payoffs. Individuals then choose behaviors to maximize their utilities or preferences given others’ behaviors. These behavioral choices in turn lead to material payoffs, and the traits affecting the utility functions evolve according to these material payoffs. If individuals do not know whom they are interacting with (and therefore cannot distinguish between opponents with different preferences), evolutionarily stable preferences coincide with individual material payoffs (Ely 2001) or a linear combination of own and others’ payoffs when there is assortment (Alger and Weibull 2013). In many cases, however, individuals can respond to others with different preferences because the behavioral game will be played repeatedly over time allowing individuals to indirectly learn each others’ preferences. In such a case, prosocial preferences can evolve to stabilize cooperation by generating positive behavioral feedbacks between individuals (Akçay et al. 2009). Importantly, this behavioral feedback acts synergisticaly with genetic relatedness in sustaining cooperation (Akçay and Van Cleve 2012; Van Cleve and Akçay 2014).

Internalized social norms can thus be seen as an evolved component of individual preferences that bias, but not necessarily dictate, individual behavior (Gintis, Helbing, and others 2015; Gavrilets and Richerson 2017). Here, we use the theoretical framework for preference evolution to ask when and how much norm internalization will be selected for, and how the presence of external punishment affects the coevolution of norm internalization and the social norm itself. Specifically, we model internalization as a subjective disutility experienced by individuals who deviate from the norm. In this setting, whether internalization evolves or not turns out to depend on whether this disutility is an accelerating or decelerating function of the deviation from the social norm. We show that in the absence of external punishment internal punishment functions that decelerate can evolve whereas internal punishment functions that accelerate cannot. When external punishment is present however, accelerating internal punishment functions yield stronger norms and more cooperation than decelerating ones. These results highlight the important role that the proximate cognitive and psychological mechanisms play in shaping whether and how norm internalization evolves.

## Modeling Framework

### External Enforcement Only

We model a population composed of groups of *n* individuals that play a public goods game with the possibility of punishment. In the first model, each individual is endowed with two traits: a normative “opinion” about what individuals in the group should contribute to the public good, denoted by *α*_*i*_, and an investment *p*_*i*_ into a punishment pool 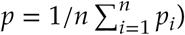, which determines the amount of punishment that can be inflicted on individuals who deviate from the norm of the group. The norm of the group, denoted by *α*, is a function of the opinions: individual opinions *α*_*i*_. For instance, we can imagine that the group norm *α* is simply the average of individual opinions:

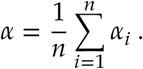

We assume individuals first make a one-time contribution to the pool that will mete out punishment for deviations from the group norm and then play a public goods game with each other. Individuals in the public goods stage follow a behavioral dynamic where they adjust their behaviors given their preferences and the behaviors of their groupmates (Akçay et al. 2009; Akçay and Van Cleve 2012). Specifically, in our first model we assume that a focal individual chooses its action, *a*_*i*_, to maximize its own payoff, denoted by *u*_*i*_ and given by:

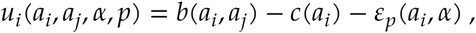

where the first term is the public good benefit from the contribution of the focal individual *a*_*i*_ and the the contributions of other individuals in the group *a*_*j*_, the second term is the private cost of contributing, and the last term *ε*_*p*_ represents the material punishment inflicted to the focal individual due to its deviation from the social norm and the mean level of punishment *p* where *ε*_*p*_(*a*_*i*_, *α*) is an increasing function of the absolute deviation |*a*_*i*_ − *α*|. Note that this payoff function reflects only the gain from the public goods stage, and thus does not include the cost of contribution to the punishment pool, reflecting our assumption that individuals make this contribution before the public goods game starts, hence it is a sunk cost at that point. We assume that all individuals adjust their behaviors dynamically until they reach a behavioral equilibrium, which happens at a sufficiently fast time-scale such that their total payoff from the public goods game is given by the equilibrium contribution levels, which we denote with an asterisk.

The payoff of a focal individual at the end of the public goods game minus the cost of the punishment contribution is the fitness of that individual. For a focal individual with normative opinion *α*_*i*_ and punishment investment *p*_*i*_ in a group with opinion *α*_*j*_ and punishment investment *p*_*j*_, the fitness *w*_*i*_ is given by

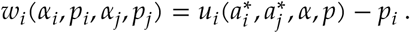

where the fitness cost of a unit of punishment is assumed to be unity. Below, we proceed to analyze this model. We first characterize the behavioral equilibrium of the public goods game given a punishment pool, then derive the first order conditions for the evolutionary stability of the normative opinion *α* and the punishment contribution *p*.

### The Behavioral Equilibrium

The first-order condition for the (monomorphic) behavioral equilibrium is given by:

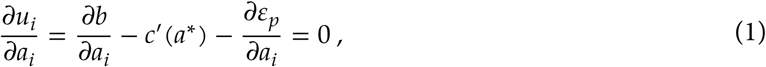

where all the partial derivatives are evaluated at 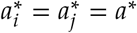. From this condition, we can read a couple things.First, assuming that the benefit function is decelerating (*∂*^2^*b*/*∂a*^2^ < 0) and the cost function is accelerating (*c*”(*a*) > 0), increasing the punishment pool *p* will have the effect of increasing the equilibrium contributions. Second, the equilibrium contribution level *a*\* will only exactly match the normative expectation *α* when the latter is equal to the individually optimal or “selfish” contribution level, which occurs when the marginal benefit equals the marginal cost in the absence of any punishment. This can be seen from the fact that when *a*\* = *α*, the third term vanishes, and the behavioral equilibrium conditions reduce to the equilibrium condition without punishment,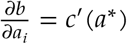. Third, any equilibrium contribution level greater than the purely selfish one can only occur when the contribution norm is at an even higher level or *α* > *a*\*. The first two terms in equation (1), will be negative since *a*\* is above the selfish level and benefits decelerate and costs accelerate. When *α* > *a*\*, the third term in equation (1), which measures the effect of a change in punishment on payoff, will be positive and can cancel the first two terms since increasing *a*\* closer to *α* decreases punishment. If the opposite is true and *α* < *a*\*, the third term will be negative since increasing *a*\* further from *α* increases punishment. In other words, if the social norm exceeds that of individually optimal behavior, players will shade their contributions to the public good to be somewhere between the individually optimal level and the normative prescription.

The behavioral equilibrium *a*\* given by solving equation (1) is a stable rest point of the behavioral dynamics whenever (Akçay and Van Cleve 2012 Appendix A3):

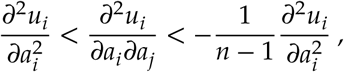

which translates to:

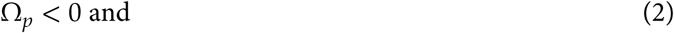

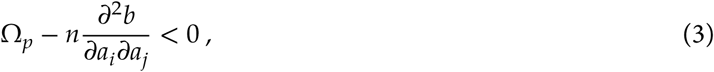

where 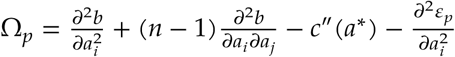

### First order ESS conditions

We can write the first order ESS conditions with population structure as in Akçay and Van Cleve (2012). The first order conditions for *p* and *α* are:

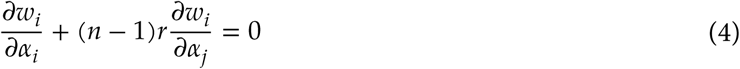

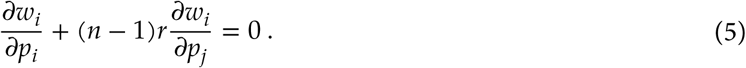

Working out the partial derivatives in equation (4) first, we have:

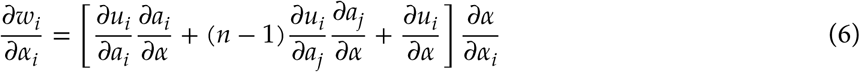

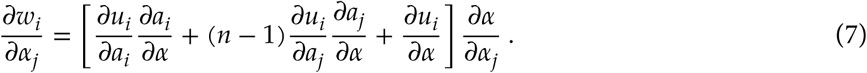

The only difference between the two equations are the partial derivatives 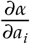 and 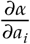 at the end. Since we are considering homogenous groups, we can assume each individual’s normative opinions has the same effect on the group norm; in other words: 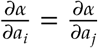, and therefore 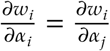. This means that equation (4) can be written as:

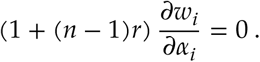

This condition implies that the evolutionarily stable contribution norm will maximize a focal individual’s fitness, taking into account the behavioral responses of the whole group to the contribution norm. Expanding the partial derivatives in equation (6) using the definition of *u*_*i*_ and using the fact that 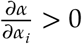, we can write the first order ESS condition for the group norm as follows:

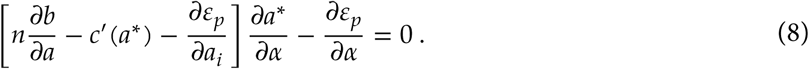

This equation yields some immediate insights: the first term on the left-hand side is positive (because it is equal to the LHS of equation (1) plus 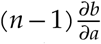, which is positive, and contributions increase with increasing *α*). This means that at ESS, the second term has to be negative, i.e., 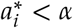, meaning that individuals underinvest relative to the normative expectation of the group. That implies that individuals are experiencing some punishment at ESS.

Likewise, for the punishment investment, *p* we can write:

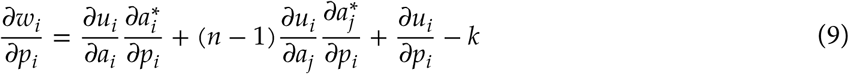

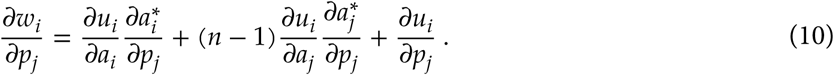

The derivatives with respect to *p*_*i*_ and *p*_*j*_ in equations (9) and (10) evaluate to 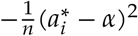. Thus, we can write the first order ESS conditions for *p* as:

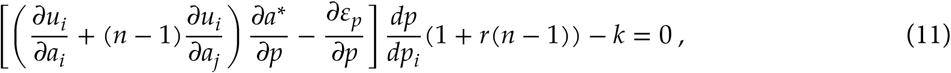

where for equation (11), we used the fact that both the focal and non-focal individuals’ punishment contribution go to a common pool that affects each individual in the same way, and hence each individual’s equilibrium contribution reacts to a change in any individual’s punishment contribution in the same way (i.e.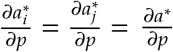)

To calculate the partial derivatives of 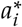 and 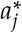 with respect to *α*, and *p*, we differentiate the behavioral equilibrium condition equation (1) with respect to the evolving variables, and solve for the relevant partial derivatives to obtain:

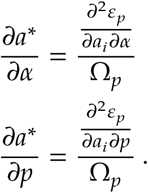

### With Internalized Punishment

In this section, we model the evolution of a psychological mechanism for internalizing norms in the presence of (fixed) external punishment. We operationalize internalization as an inherent motivation to follow the prescribed behavior. Specifically, we assume agents have an evolving trait,*τ*, that determines how much internalized “discomfort” they feel due to deviations from the prescribed norm. Mathematically, we assume now that our agents maximize the following objective function during the behavioral dynamics:

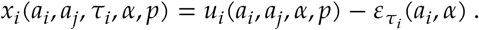

where 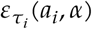 is an increasing function of the deviation from the norm, *a*_*i*_ -*α*. The 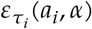 term is analogous to the external punishment term included in the payoff *u*_*i*_, but is only felt subjectively, with no direct effect on the material payoff of the individuals. However, the presence of such subjective discomfort can alter the behavior of the focal individual and can therefore have an indirect effect on its payoff.

The behavioral equilibrium condition for a focal individual is now given by:

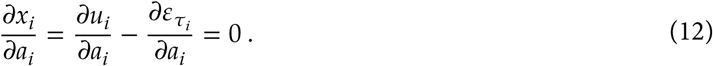

This condition implies that the effect of internalized discomfort (positive *τ*) is similar to external punishment: as long as the group norm *α* is higher than the individually optimal contribution maximizing *u*_*i*_, the effect of *τ* is to increase contributions. The stability of the behavioral equilibrium is again determined by the same conditions as equations (2) and (3), except Ω_*p*_ is replaced by 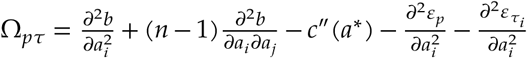.

The partial derivatives of the equilibrium contribution *a*\* with respect to *α* and *τ* are given by:

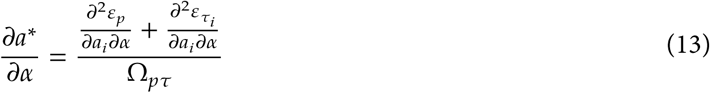

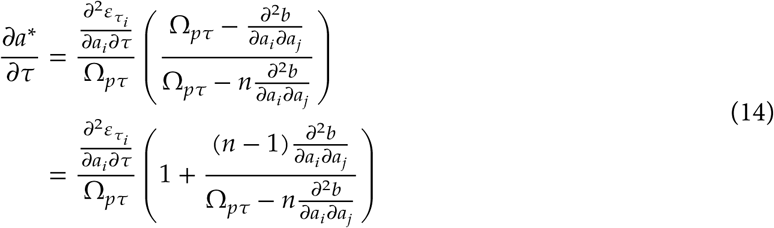

The ESS condition for *τ* is given by

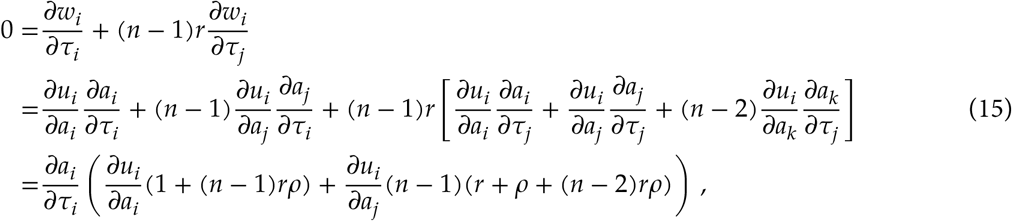

where 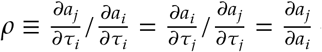 denotes the responsiveness of an individual’s contribution to the change of another individual’s contribution (Akçay et al. 2009; Akçay and Van Cleve 2012). At the monomorphic equilibrium, the responsiveness of any individual to any other individual is the same, which allow us to write the last line in equation (15). So long as *a*_*i*_ ≠ *α*, then 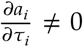, and satisfying the first order ESS condition requires that the term in the parentheses in equation (15) has to vanish. By setting this term to zero and rearranging we obtain

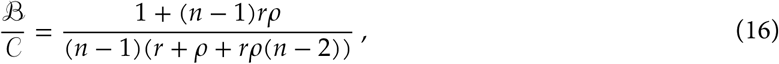

where the marginal benefit to the focal individual from the investments of others is 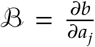 and the net marginal cost from its own investment is 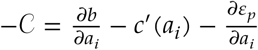. Because *τ* does not affect the payoffs of individuals directly, the response coefficient *ρ* can be written as in Akçay and Van Cleve (2012 eq. 13):

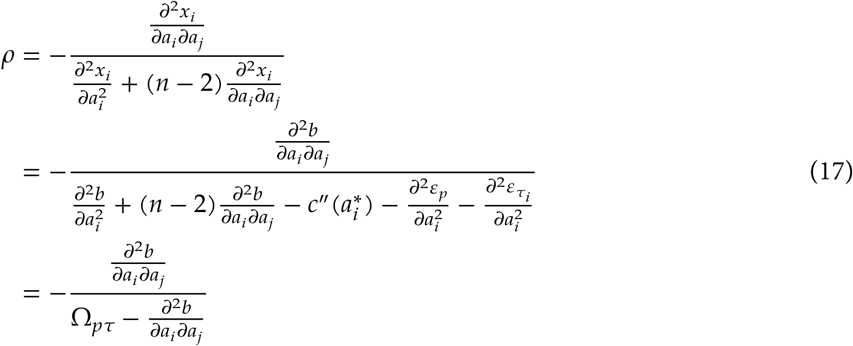

Further, from the behavioral equilibrium conditions (A27) in Akçay and Van Cleve (2012), the response coefficient *ρ* must satisfy the following inequality: −1/(*n* - 1) < *ρ* < 1.

Since *τ* does not affect the payoffs directly, the ESS condition for *α* stays the same as equation (8) except that all derivatives are evaluated at the behavioral equilibrium solving equation (12).

### Representative Functions

In order to numerically analyze the model, we need to specify a few particular functions for the cost, benefit, and punishment functions. These functions are meant to capture our intuitions about public goods payoffs, and different notions of how external and internalized enforcement of norms might work.

First, we introduce a public good benefit function *b*:

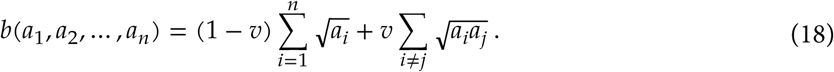

This function captures two important intuitions about public goods benefits. First, the fact that all contributions are in square roots ensures that there will be (eventually) diminishing results from contributions to the public good so that a finite contribution is socially optimal. Second, it allows individual contributions to be synergistic or anti-synergistic (or complements or substitutes in economic terminology), as modulated by the parameter *v*, which we assume is in the range [-1, 1]. Specifically, the second sum of equation (18) corresponds to individuals’ contributions interacting pairwise. Positive *v* can be interpreted as representing collaborative interactions that contribute positively to the public good. Negative *v* on the other hand, represents agonistic or competitive interactions that diminish total public goods provision. Thus, this function allows us to represent a range of social scenarios.

For the cost of contribution to the public good, *c*(*a*), we use a simple quadratic function that represents accelerating marginal costs:

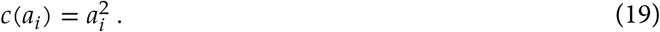

The external and internalized punishment functions, *ε*_*p*_(*a*_*i*_, *α*) and 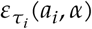, respectively, both increase when the deviation of a focal individual’s contribution from the group norm |*a*_*i*_-*α*| increases. However, the behavioral and evolutionary stability conditions above also depend on the curvatures or second-order derivatives of these functions. For external punishment,*ε*_*p*_ (*a*_*i*_ *α*) we use the following functional form:

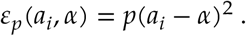

This function captures the notion of graduated punishment (Ostrom 1990) as applied to our setting, where small deviations from the norm encounter relatively small punishments, but the punishment escalates with larger deviations. The size of the punishment pool, *p*, modulates the amount of punishment. For internalized punishment function, we investigate two forms with different curvatures. The first form is analogous to the external punishment function above and is accelerating in terms of the deviation of contribution from the norm:

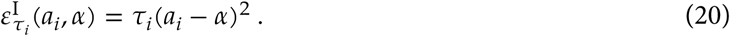

The second form captures the notion that as an individual’s deviation from a norm grows, they might not experience infinitely increasing discomfort. Rather, individuals that are already far from a norm may feel relatively small additional discomfort for the same additional deviation compared to individuals who are closely adhering to the norm. This suggests an internalized discomfort function that plateaus at large deviations from the norm. We use the following function to represent this case:

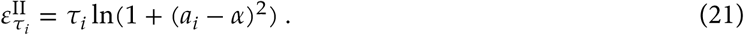

This function behaves similar to 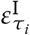 at small deviations from the norm, but saturates at large deviations.

## Analysis and Numerical Results

### External Punishment Only

First, we assume that only external punishment is possible. Figure 1 depicts the evolved contribution and punishment levels and the evolved norm using the benefit and cost functions in equations (18) and (19) with no synergistic interactions or *v* = 0. It shows that an evolutionarily stable (ES) social norm *α*\* and external punishment level *p*\* need a threshold level of relatedness within social groups. Below this threshold value of relatedness, a positive contribution to the punishment pool is not evolutionarily stable, which reflects the second-order dilemma that costly punishment poses. As relatedness crosses a threshold value, the ES punishment level increases from zero, positive selection arises on the social norm *α*, and an ES *α*\* and contribution level evolve. With increasing relatedness, punishment contributions increase while the contribution norm decreases. This is due to the fact that more relatedness allows for higher *p* and more punishment induced cooperation. However, higher *p* increases the marginal cost of maintaining a social norm at a particular level, and this cost is increasingly paid by relatives. Despite decreasing the social norm *α*\*, increasing relatedness increases the equilibrium contribution to the public good (*a*\*) and the net fitness *w*. Thus, as relatedness increases, groups become more cooperative because they punish smaller deviations from the norm more harshly.

**Figure 1:**
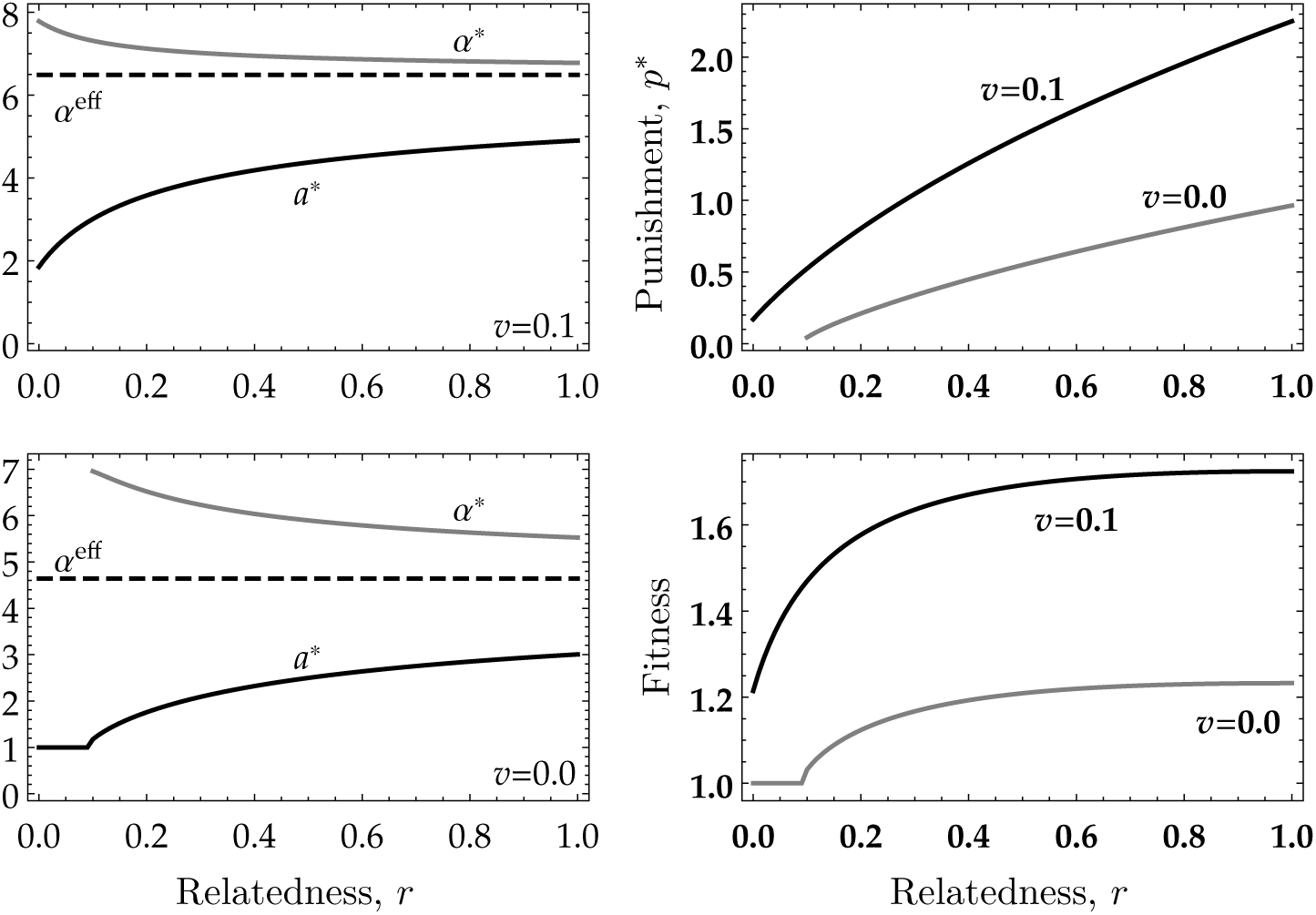
The evolutionarily stable (ES) contribution norm *α*\*, punishment investment *p*\*, and fitness as a function of relatedness for no complementarity, *v* = 0.0, and positive complementarity, *v* = 0.1 given *n* = 10 individuals. The norm and investment levels in the first column are relative to the baseline investment (ES investment and norm with no punishment). The fitness values in the lower right panel are relative to the baseline fitness (ES investment and norm with no punishment). For low values of relatedness and *v* = 0.0, no positive punishment investment is evolutionarily stable, so all norms are neutral, since they do not get enforced.

Figure 2 further shows that the critical value of relatedness needed to sustain a normative equilibrium decreases with increasing complementarity of contributions to the public goods. This is intuitively due to the fact that when contributions are complementary, a coordinated increase in the contribution level (actions) of multiple individuals generates more benefits for everyone than if contribution levels are substitutes. Therefore, external punishment, which has the effect of increasing everyone’s contribution, has a higher benefit in situations where actions are complementary.

**Figure 2:**
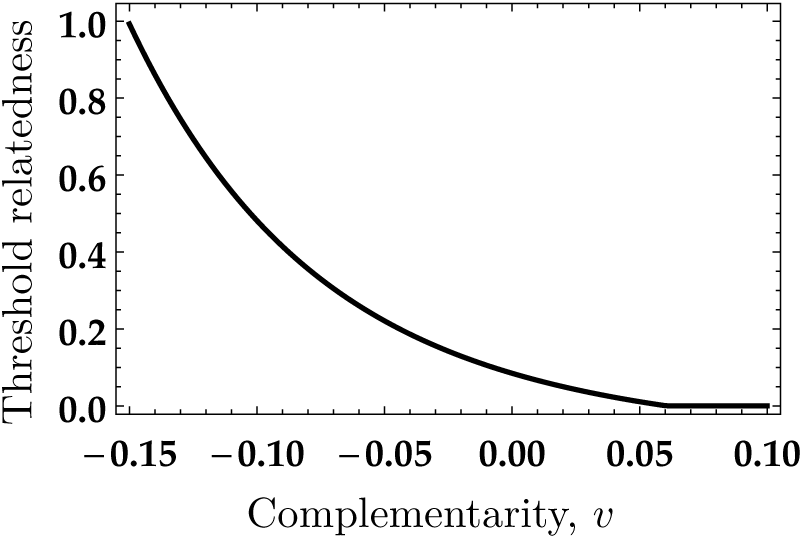
The critical value of relatedness needed for a normative equilibrium to be ES as a function of the complementarity parameter *v*. As the public good contributions get more complementary (or individuals become more collaborative), lower relatedness is needed to sustain a public goods contribution norm.

### Internalized Punishment Only

Now, suppose that there is no external punishment, *p* = 0, but internal punishment *τ* can evolve. We show in this case that norm internalization can only be evolutionarily stable if the internalized punishment function becomes saturating at some point. When there is no external punishment, the ESS condition for the contribution norm *a*, equation (8), reduces to:

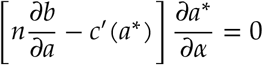

where 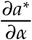 is given by eq. equation (13). This means that the first order ESS condition for *α* can only be satisfied if either 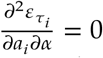 or the term in the square brackets vanish. The latter implies that the behavioral equilibrium maximizes total payoff of the group (Akçay and Van Cleve 2012), which in turn means that the left hand side of the ESS condition for *τ*, equation (16), equals 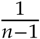. Satisfying the ESS condition for *τ* then requires either *ρ* = 1*or r* = 1 (or both). The clonal condition with *r* = 1is an uninteresting case, since in clonal groups no conflict over the public good exists. The other option, *ρ* = 1from equation (17) implie s that. Ω_0*τ*_ = 0, but this contradicts the stability condition for the behavioral equilibrium (equation (2)). This means that for a social norm to exist with purely internalized punishment,

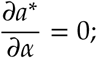

i.e., the internalized enforcement of the norm should be such that at some point, increasing the group norm does not yield high er contributi on. In other words, a necessary condition for an ESS with an internalized punishment only is that at some point increasing the social norm does not elicit higher contributions from group members. The intuition behind this result is that traits that raise the group norm *α* are not costly without norm enforcement (internal or external) but in the presence of norm internalization, they elicit higher contribution s from group-mates. Thus, as long as 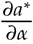 is positive, selection will act to increase the group norm. Only when norm internalizers stop responding to higher *α* values can we satisfy the ESS conditions.

Using equation (13) in the absence of external punishment, we can see that 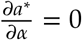 requires

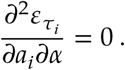

This condition means that the internal punishment needs decelerate, or start to saturate, at high deviations from the social norm. In other words, if individuals are already showing a big enough deviation from the norm, increasing the norm will not necessarily impose a high enough marginal discomfort on them to make them increase their contribution. In colloquial terms, these individuals would be “giving up” on trying to keep up their contribution level with the norm. Without external punishment, the contribution norm will evolve precisely to the level at which individuals are giving up on following it. This can lead to relatively high levels of cooperation, as all individuals will respond with the same increase in contribution level with an increase in the norm (until they stop responding), meaning that the marginal cost to the focal individual of contribution more will be offset by the equivalent contributions from group-mates.

### Internalization in the presence of external punishment

Finally, assume that there is a fixed level external punishment and that the level of social norm internalization *τ* can evolve. Figures 3 and 4 depict the ES contribution norm and internalization level as a function of relatedness and external punishment under different internal punishment functions. For both accelerating and decelerating internal punishment functions, higher relatedness results in higher internalization. For accelerating internal punishment (Figure 3), a minimum level of relatedness has to be present before internalization, *τ* > 0, can evolve (these low relatedness values are not plotted in Figure 3). For accelerating punishment, the ES contribution norm decreases with increased external punishment: mirroring the case with external punishment only, increases in external punishment increase the cost of maintaining a high social norm, which generates selection for a lower contribution norm. Increases in the degree of complementary or synergy of the benefit of contribution (*v*) increase the ES norm and contribution level but have little effect on the level of internalization.

**Figure 3:**
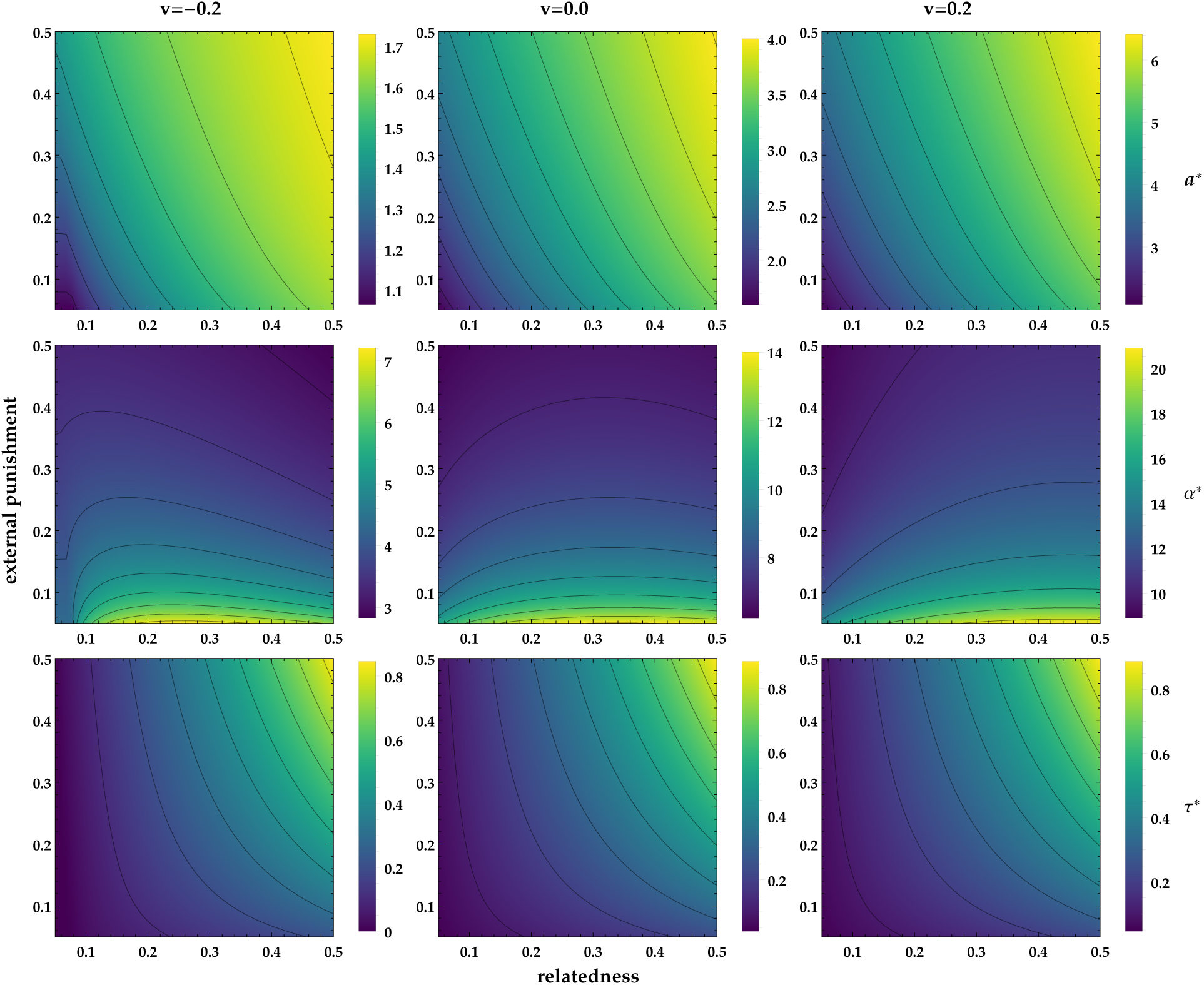
The ES contribution *a*\*(top row), ES norm *α*\* (middle row), and ES internalization *τ*\* (bottom row) for an accelerating internal punishment function (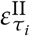, in equation (20)). Columns represent benefit functions varying from antisynergistic to synergistic going left to right. The ES contribution level *a*\* and norm *α*\* are shown relative to their values with no punishment (external or internal). The norm decreases with *p* and increases with *r* monotonically under all benefit functions, while norm internalization increases with both. The resulting contributions to the public good increases with both external punishment and relatedness despite the fact that the contribution norm might decline with external punishment.

**Figure 4:**
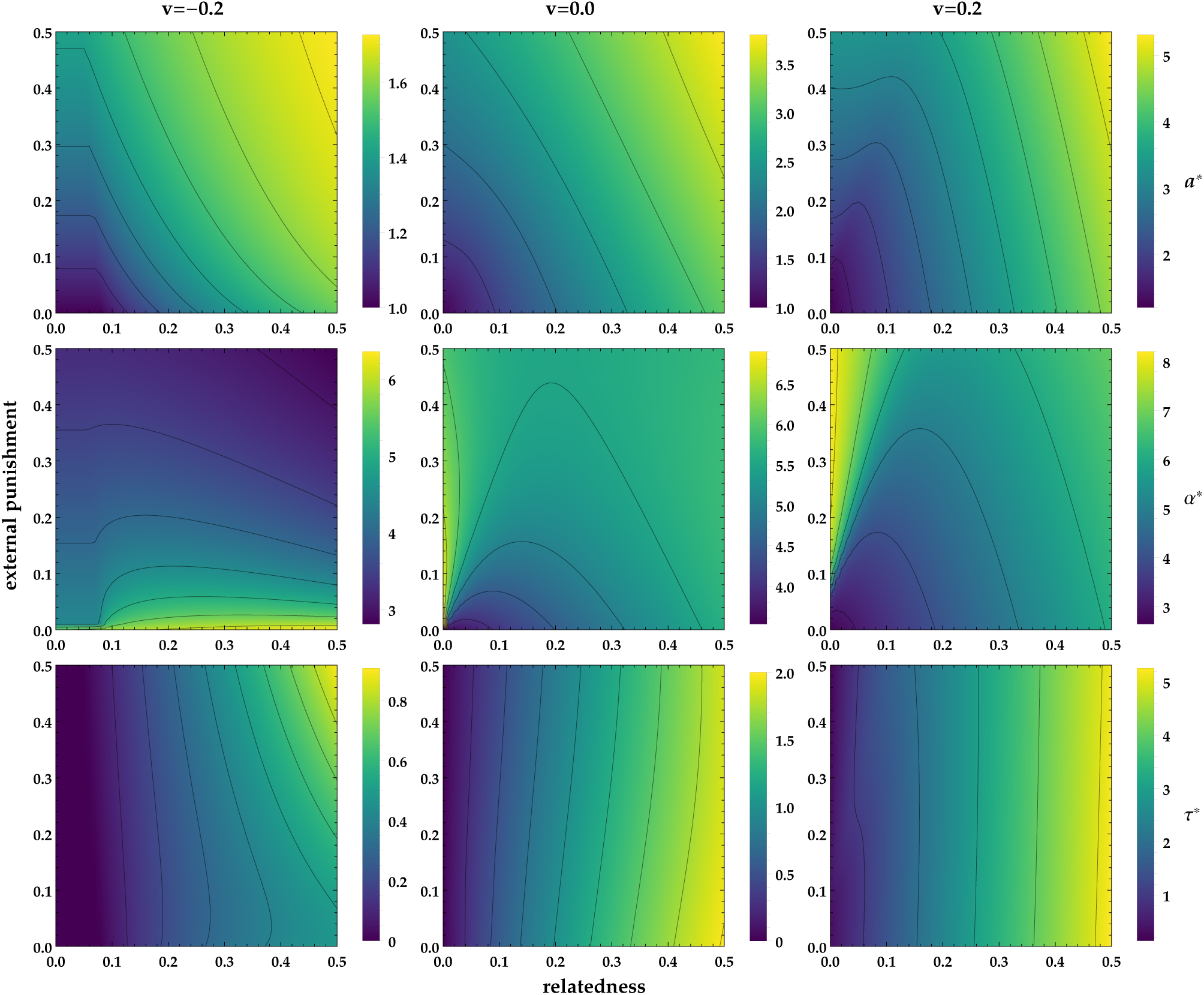
The ES contribution *a*\* (top row), ES norm *α*\* (middle row), and ES internalization *τ*\* (bottom row) as a function of relatedness and exogenously fixed external punishment, with decelerating internal punishment (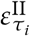, equation (21)). The ES contribution level *a*\* and norm *α*\* are shown relative to their values with no punishment (external or internal). In the dark region for *v* = −0.2, the ES *τ\** = 0, so the social norm is purely maintained by external punishment. For antisynergistic benefits, increasing external punishment decreases the contribution norm, while for additive and synergistic benefits, it increases the norm. In all cases, norm internalization increases with relatedness but is relatively unresponsive to external punishment. The ES contribution generally increases with both external punishment and relatedness, but remains at lower levels compared to the accelerating function in Figure 3.

However, Figure 4 shows that decelerating internal punishment produces more complex patterns. When the contributions to the public goods are antisynergistic (*v* < 0), the patterns mirror those with accelerating internal punishment: the ES contribution norm decreases with increases in external punishment and a threshold relatedness is required for the ES *τ* to be positive. In contrast, when the benefit function is additive or synergistic (middle and right columns of Figure 4 where *v* = 0 and *v* > 0, respectively), the ES contribution norm increases with increasing external punishment but changes non-monotonically with changes in relatedness: *α*\* decreases at low relatedness and increases at higher relatedness. Furthermore, the level of the ES contribution norm and resulting ES contributions (relative to their levels with no punishment at all) are significantly lower for the decelerating internal punishment function as compared to the accelerating one.

These results, together with the analytical result above for the case of no external punishment, suggest a trade-off. While decelerating internal punishment functions might be easier to evolve in the absence of external enforcement, they might be less efficient in maintaining high cooperation than accelerating internal punishment, once the latter is stabilized by external enforcement.

## Discussion

Not only have humans evolved a capacity to follow social norms, they can also internalize those norms so that they are intrinsically motivated to comply with group- or population-level standards of behavior. Such internalization is frequently crucial to the proper functioning of human groups and society. We have presented a model for how the capacity for internalization might evolve through evolution of social preferences. In particular, our model explores how external enforcement in the form of punishing deviations from the norm affects the evolution of norm internalization in the form of a preference for following the norm or a subjective discomfort from failing to follow the norm.

A main result from our model is that the interaction between external punishment of norm deviations and the evolution of norm internalization crucially depends on the shape of the function that describes the discomfort or internal punishment experienced by individuals. We distinguish between two functional forms for this internal punishment function: (i) accelerating functions representing cases where the marginal discomfort from deviation keeps increasing the farther an individual is from the norm, and (ii) decelerating functions, where at large enough deviations from the norm, the marginal discomfort starts decreasing and may vanish (i.e., individuals do not feel *additional* discomfort from deviating more). When internal punishment is an accelerating function, the evolution of internalization requires the presence of external punishment, while with a decelerating internal punishment function, internalization can evolve even if there is no external punishment. Accelerating internal punishment is not stable on its own because it causes a focal individual to always respond to an increase in the group norm by increasing their contributions. That means that groupmates of this focal individual could increase the value of the norm *α* and elicit more contribution from the focal individual. In the absence of external punishment, there is little cost to an individual increasing their part of the contribution norm *α*, but there is a significant benefit since everyone will contribute more. This causes the social norm to increase beyond optimal, and in response, the level of internalization *τ* is selected to decrease. This process eventually drives *τ* down to zero even as the norm *α* evolves to large values that are not enforced. This run-away increase in the norm and decrease in the internalization is thwarted when there is external punishment, since increasing the norm is now also directly costly to a focal individual due to the increased punishment. The runaway dynamics are also short-circuited when the internal punishment function starts leveling off, because as the norm keeps increasing, even an internalizing focal individual stops feeling the necessary additional discomfort to try to keep up. That caps how much the contribution level of a focal individual will increase in response groupmates increasing the norm and stops the norm from increasing too much. This contrast between the two functions continues in the presence of external punishment. For accelerating functions, higher external punishment increases the strength of internalization (Figure 3), while with decelerating functions internalization is most responsive to relatedness between group members, as opposed to external punishment (Figure 4).

These results suggest that internal punishment that eventually levels off might be more robust for evolving norm internalization on its own and that evolved norm psychology should have a mechanism for capping the subjective discomfort from norm violations. When this happens, individuals do not experience large marginal punishment or reward from marginal changes to their behavior or the norm. This bears some resemblance to the notion of goal disengagement (Wrosch et al. 2003) in psychology, which describes situations where individuals stop pursuing unattainable goals. In our model, the disengagement is “local” in the sense that individuals do not completely give up on prosocial behavior, but prove unwilling to go beyond a certain level of contribution. In a heterogeneous population (e.g. with individuals with different endowments), the point at which individuals disengage would vary, which might put pressure on group norms to diversify, leading to socially stratified social norms.

Further, our results show that presence of external punishment can drive the evolution of internalization. This effect is most pronounced for accelerating internal punishment functions, where external punishment keeps the social norm from increasing too much due to the fitness costs stemming from such punishment. With the social norm constrained by external punishment, internalization can evolve to take over some of the enforcement function, and reduce the amount of material punishment at equilibrium. In contrast, with decelerating internal punishment, internalization is much less sensitive to external punishment. In both cases, norm internalization becomes stronger with increased relatedness in a population. This is not surprising, since internalization is a trait directly benefiting group mates, so is expected to be favored when relatedness between group members is high. Under decelerating costs of internal punishment, but not under accelerating costs, the level of internalization also increases with the degree to which contributions to the public good are synergistic. However, since accelerating costs produce much stronger discomfort at the same level of internalization, *τ* accelerating costs still produce greater stable contribution levels and norms. Thus, if external punishment is available through some social institution, accelerating internal punishment may produce more powerful norms with greater levels of cooperation.

There are only a handful of previous models of how the capacity to internalize social norms might have evolved. In an influential paper, Gintis (2003) developed a gene-culture coevolution model where a genetically transmitted allele allows for the social acquisition and internalization of a norm from parents and others. He found that such an allele will fix in a population if it allows the acquisition of individually beneficial norms, and that altruistic norms can “hitchhike” on this internalization capacity to also establish. In our model, internalization does not hitchhike on another beneficial trait; instead, it evolves because it allows individuals to reduce the punishment they experience and it generates positive behavioral responses. Another difference is that we consider both norms and the internalization to be inherited by the same mechanism (genetic or cultural), whereas Gintis models the co-evolution of traits transmitted genetically and culturally. Considering gene-culture coevolution in a setting like ours will be an interesting future direction. More recently, Gavrilets and Richerson (2017) provide agent-based simulations of a setup closely related to ours, where individuals can contribute to a public good in a group as well as to the effort to punish deviations. Similar to our paper, they model internalization as an intrinsic reward for contributing to the public good and punishment, with a linear reward function and a fixed contribution norm. They find that internalization can evolve more easily in games where groups compete only indirectly (i.e., group success depends only on its own public good production; “us-vs-nature” games), compared to when groups compete directly (where group success depends on other groups’ production; “us-vs-them” games). In the latter case, selection already favors high amounts of investment and therefore internalization is not required to sustain cooperation. Interestingly, Gavrilets and Richerson (2017) find that under many parameter regimes, substantial genetic variation in norm internalization is maintained, including dimorphisms between high norm internalizers and non-internalizers. Our analysis did not explicitly look for diversifying or balancing selection that could produce such polymorphisms, which is another interesting question for future research.

## Conclusion

The phenomenon of internalization in humans is complex and is affected by many different processes at the individual, group, and societal levels. Here, we focused on the selective pressures that come into play for the coevolution of social norms and the intrinsic motivations to adhere to them in a public goods game setting. We find that the evolution of such intrinsic motivations is predicated on how the underlying mechanism processes deviations from the social norm and encodes the discomfort, guilt, or internal punishment from falling short. We find that internalization requires external punishment when the internal punishment is accelerating whereas functions that eventually level off in the deviation from the norm can support social norms through purely intrinsic preferences. Crucially, norm internalization can create a new conflict by generating an incentive to keep increasing the contribution norm as in the accelerating case or remove conflicts by causing coordinated increases in contributions in the decelerating function case. These conflicts or their resolution will reciprocally interact with the dynamics of institutions, social networks, and other group-level processes. Understanding these interactions remains an important goal of social evolutionary theory.

## Acknowledgements

We thank W. Wilczynski, S. Brosnan, H. Gintis, and B. Morsky for comments that improved the manuscript. EA acknowledges support from Army Research Office (W911NF-17-1-0017) and Defense Advanced Research Projects Agency Next Generation Social Science Program (Grant D17AC00005). JVC acknowledges support from National Science Foundation (Grant #1846260).

